# Assessing a commercial capillary electrophoresis interface (ZipChip) for shotgun proteomi applications

**DOI:** 10.1101/559591

**Authors:** Aimee Rinas, Conor Jenkins, Ben Orsburn

**Author notes:** Contributed equally to this study (Aimee Rinas, Conor Jenkins).

## Abstract

Capillary electrophoresis coupled electrospray ionization mass spectrometry (CE-MS) has long existed as a theoretical alternative to liquid chromatography coupled mass spectrometry (LC-MS). Until recently, however, the coupling of these technologies has occupied only a small niche within specific applications. A recent innovation in CE-MS is the ZipChip interface system from 908 devices that was pioneered by the Ramsay lab at NC State. This newly available source offers advantages over previous CE-MS interfaces including both relative ease of use and direct compatibility to thousands of mass spectrometers currently in use throughout the world with no hardware alterations. The ZipChip CE-MS has been demonstrated in recent studies to provide high resolution and rapid separations for the analysis of intact proteins, glycoproteins and glycosylated peptides, with more applications likely on the way.

In this study we assess the capabilities of the ZipChip system in the context of high throughput global shotgun proteomics experiments. We find that on a high field Orbitrap system we can repeatedly identify as many as 800 unique protein groups in an experiment using a run time of 12 minutes. We find the ZipChip CE-MS system to be widely applicable for both data dependent and data independent acquisition experiments as well as targeted experiments. We conclude that the ZipChip is an attractive alternative solution to traditional nanoflow ESI-MS/MS for the analysis of the genomes of single celled organisms and for offline fractionation of eukaryotic proteomes.

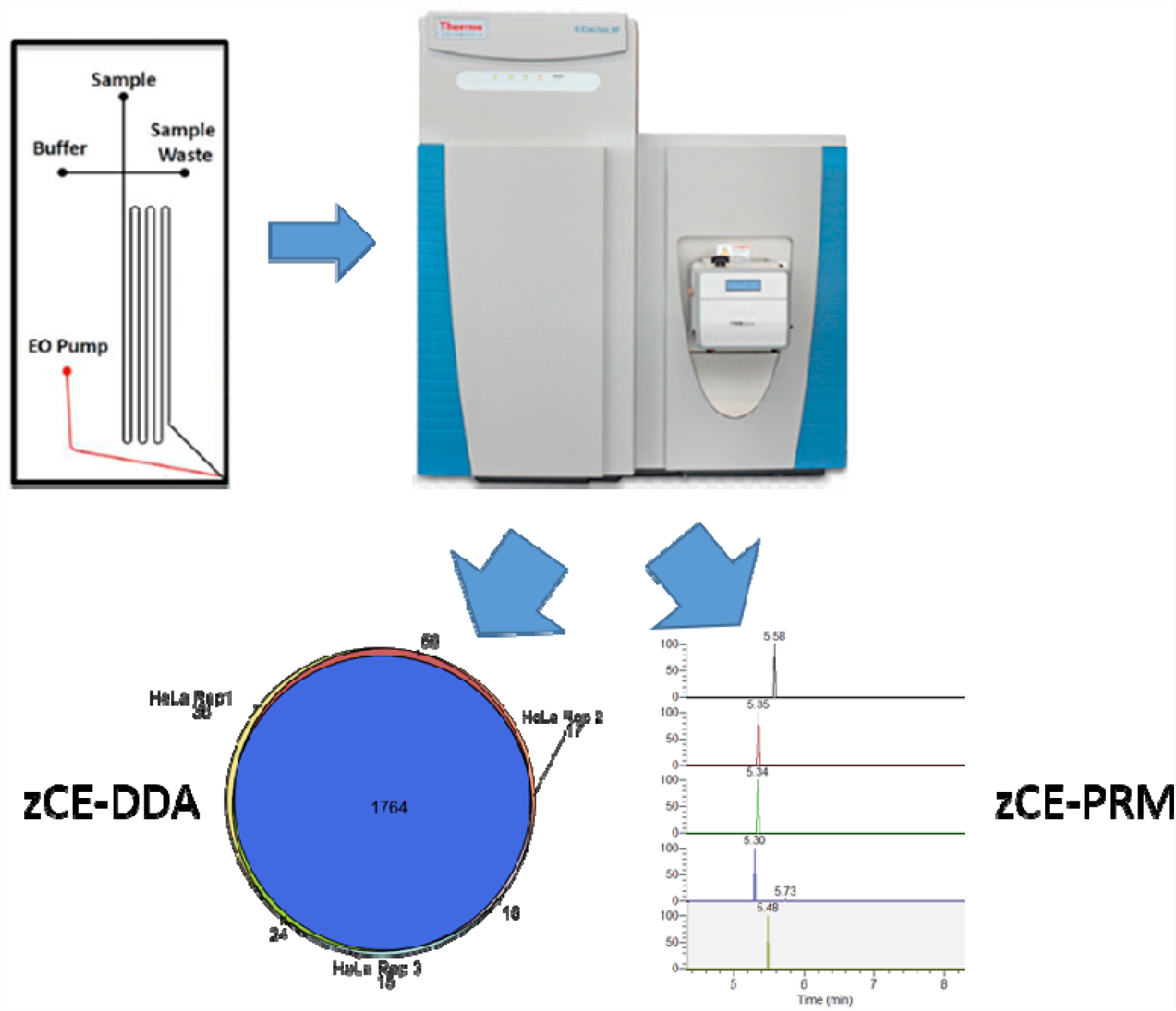

## Introduction

Since the widespread adoption of nanoflow HPLC (nLC) coupled mass spectrometry for shotgun proteomics, which we will refer to here as nLC-MS, the technique has become nearly ubiquitous. Currently, the majority of shotgun proteomics systems consist of a nanoLC separation system coupled to a high-or ultra-high resolution mass spectrometer linked by electrospray ionization^1^. While the capabilities of mass spectrometers themselves have increased at a remarkable pace since - with improvements in resolution, mass accuracy, speed and sensitivity leading the way, nLC can be considered a bottleneck for many experiments. ^2,3^Improvements in chromatography materials and the introduction of higher pressure pumps have resulted in overall improvements in the reproducibility and resolution of nLC separations. However, operating nLC systems is still not a trivial matter. Due to the high sensitivity of the systems, carryover of sample components from one run to the next remains a major concern. Reproducibility of separations are also challenging. Commercial columns that have been subjected to rigorous quality control and assurance procedures are now available from many vendors. These columns are often limited in available separation chemistries or may be prohibitively expensive for many labs, leading many labs to resort to in-house packed columns. For example, the recently initiated CPTAC 3 initiative has chosen to use in-house packed columns, presumably due to the absence of a commercially manufactured column utilizing the relatively new ReproSil-Pur C-18 1.9μm separation material.^4^

Recent work from the Qu lab at the University of Buffalo has demonstrated that reproducible nLC is achievable for both extremely large sample cohorts within single studies ^5^, as well as intra-study over large periods of time. The IonStar methodology requires rigorous attention to sample preparation ^6^ and loading, as well as the use of 100 cm columns and data processing that integrates peak features and peak integration features involved in false discovery calculations ^7^. Many labs, however, rely on low resolution offline fractionation with concatenation followed by nLC separation on columns typically 20 cm − 35cm in length. Recent work from the Olsen lab has challenged this established paradigm by utilizing high resolution high pH reversed phase fractionation and LC-MS gradients as short as 32 minutes ^8^, which was later further reduced in a follow-up study utilizing a faster mass spectrometer ^3^.

Capillary electrophoresis linked mass spectrometry (CE-MS) is an alternative solution to nLC-MS workflows. Due to the mechanism of separation, sample carryover from one run to the next may be considerably diminished compared to nLC ^9^. The resolution of CE separations may also exceed that of LC based separations and may lead to the baseline separation of many compound classes, such as glycans that can not be resolved by traditional nLC ^10^. The resolution of the CE separations has traditionally been a challenge for mass spectrometry, as high-resolution mass spectrometers have struggled to match the speed of CE compound elution ^11^.

Recent advances in proteomics LC-MS instruments have featured marked increases in scanning speed. The Q Exactive HF can achieve a realistic proteomics sequencing speed of 22 Hz ^12^ and a further advance on this platform, the HF-X can nearly double this number, albeit with lower resolution MS/MS scans ^3^ Software advances for the Orbitrap Fusion instruments in 2018 (manufacturer communication) allow an Orbitrap scan speed of nearly 30 Hz. A recent report utilizing a Fusion Lumos system with parallelization of the Orbitrap and ion trap reportedly approaching 60 Hz and allowed efficient coupling and shotgun proteomics using a CE system ^13^.

In this study we evaluate a commercially available CE-MS interface, the ZipChip™ from 908 Devices, which utilizes microchip CE technology pioneered by the Ramsay lab for separation and direct ionization ^14^. We coupled this system to a QE HF system (zCE-MS), and evaluated standard shotgun proteomics applications. We conclude that the short gradients separations CE-MS is an attractive solution for the analysis of relatively simple samples consisting of up to 1,000 unique proteins. The zCE-MS system can provide rapid single-shot analysis of the proteomes of single celled organisms possessing relatively small genomes. For affinity enrichment or purification experiments, the low sample to sample carryover and short sample acquisition time may make zCE-MS a superior system to nLC-MS. For highly fractionated complex proteomes, the ZipChip may be a suitable replacement for nLC systems for both proteomic coverage and quantitative proteomic experiments.

## Materials and Methods

### Samples

Both the retention time calibration standard (PRTC) standard and HeLa digest standard were purchased from Pierce. The PRTC mixture was diluted into the HeLa lysate digest background with a starting concentration of 200ng of HeLa and 400fmol PRTC per microliter of 0.1% formic acid. For LOD/LOQ experiments in complex background, the PRTC was 10x serial diluted with 0.1% formic acid and used to resuspend a 10x HeLa lysate digest for PRM and LOD studies. The Yeast tryptic digest from Waters was diluted in 0.1% formic acid to a starting solution of 2ug/μL.

### Operation of the ZipChip interface

The ZipChip system was operated according to manufacturer protocol for high resolution separation as previously described ^15^. Briefly, the device was positioned on the inlet of a Q Exactive HF (Thermo). The 29 cm long separation channel of ZipChip HR was used for all experiments described. Background electrolyte (BGE) was 50% Acetonitrile containing 0.1% Formic Acid. The settings of CE were as follows: Injection load time of 5s at 2 psi, analysis run time of 12 min, pressure assist start time of 10s, CE voltage 20 kV, ESI voltage 2 kV and shield voltage of 500 V.

### Mass Spectrometry Parameters

A Q Exactive HF system equipped with Protein mode and the BioPharma accessory package was used for all experiments. Neither protein nor BioPharma mode were employed in any experiment described here. For shotgun proteomics studies an MS1 scan resolution of 60,000 with an AGC target of 3e6 and maximum fill time of 50ms was followed by a “top 10” experiment with fragmentations at 15,000 resolution. The AGC target for MS/MS was 1e5 with a maximum fill time of 30ms. Prescan calculations were performed by modeling chromatography on a 3 second peak (FWHM). Ions were collected for fragmentation using a 1.4 Da isolation width and with a normalized collision energy (NCE) of 27. For PRM experiment, an MS1 scan was performed as described above. PRMs for 4 PRTC peptides were included in the targeted list using a 2.0 Da isolation width centered on the monoisotopic mass with a 100ms maximum injection time, a 2e5 AGC target and an NCE of 35. All fragment ions were read at 30,000 resolution.

### Data Processing

For global proteomic analysis all RAW files were processed using Proteome Discoverer 2.2. Briefly, the SequestHT ^16^ and MSAmanda ^17^ nodes were used with a 10ppm MS1 tolerance and 0.02 Da MS/MS cutoff. Analysis of chimeric spectra was performed with the CharmeRT workflow in Proteome Discoverer IMP 2.1, and MSAmanda 2.0^18^ HeLa samples were compared to the canonical human UniProt/SwissProt database downloaded in April 2017, parsed on “sapiens” and the cRAP contaminant database from www.GPM.org. Yeast digest was processed in the same manner, utilizing a database parsed on “*Saccharomyces cerevisiae*” Percolator was used to approximate a 0.01 FDR confidence and peptide group FDR was approximated to the same confidence level according to software defaults.

The computer hardware for all analyses was a was a custom workstation from OmicsPCs of Baltimore MD, purchased in 2013, and fitted with a custom CPU overclock optimized for the MSAmanda search engine, with a base clock speed of 4.7Ghz on all 8 cores. The PC has 16GB of dual channel RAM and all processing was performed on a Samsung Evo solid state drive set in Rapid Mode.

## Results and conclusions

### Complex mixture data dependent analysis

A 200ng/μL solution of HeLa peptide standard was repeatedly injected using a ZipChip 12minute separation program. Identical, repeat injections of 2μg/μL of yeast digest were also analyzed using identical methodology. An overlay of the base peak chromatograms in Figure 1 displays the comparison of 2 yeast digest separations.

**Figure 1.**
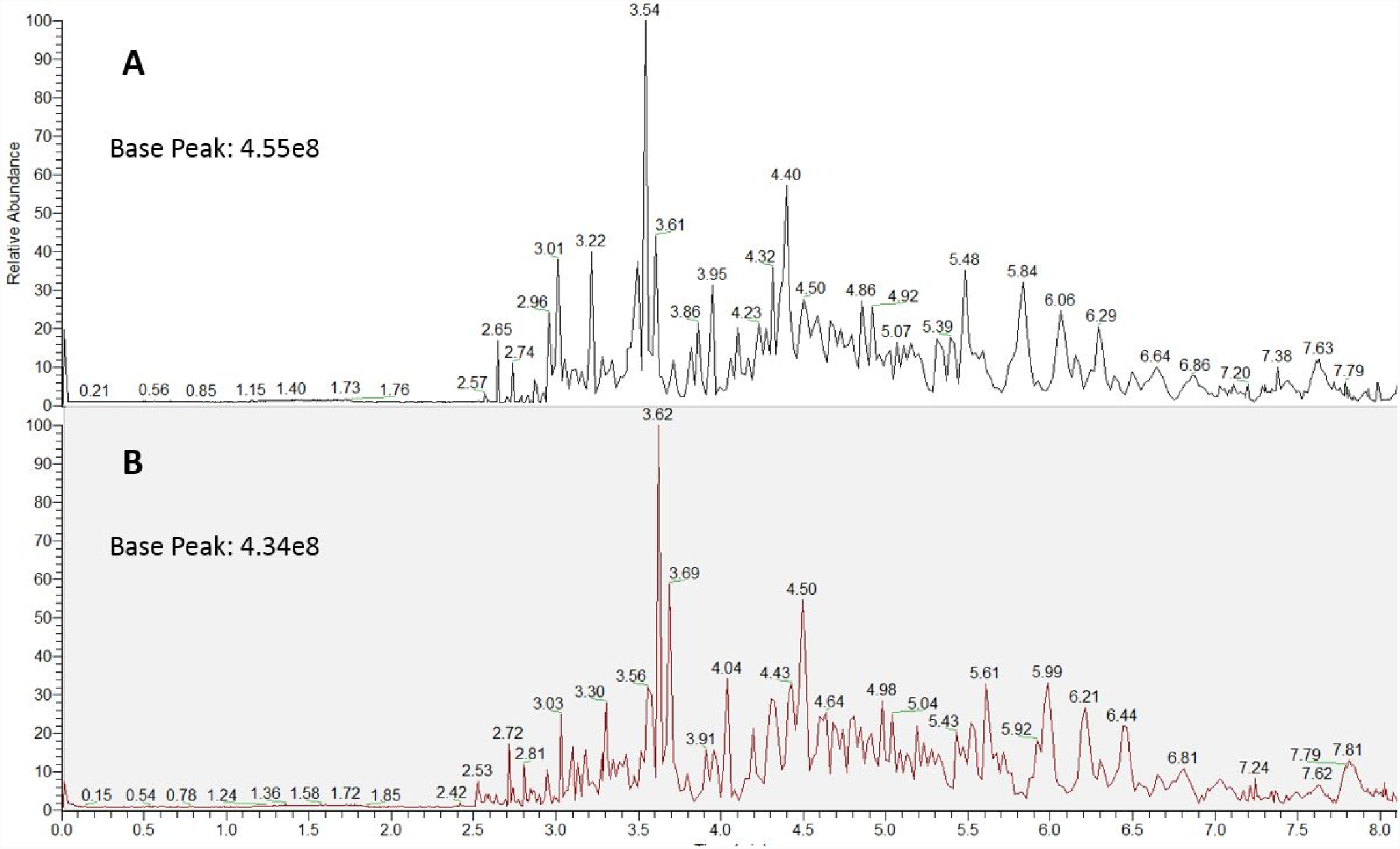
Two representative base peak extractions for yeast digest samples.

The ZipChip system allows multiple injections to be taken out of the same well without reloading the sample, despite this the base peak intensity appears to remain consistent for up to 6 injections tested. Despite expectations, the loading concentration does not appear to decrease markedly as samples are injected out of the single well for up to the 6 injections we tested. We must conclude that the ZipChip system loads only small fraction of the peptides within the well during each injection loading and suggests that preconcentration of samples may be necessary to achieve nLC levels of sensitivity. While it is challenging to equate concentrations of peptides loaded from TIC alone, a comparison of single peptides from short gradient nLC injections of HeLa suggests that the true load per run may approximate as little as 50ng of peptides per injection (data not shown). Despite the low intensity from these loads each run resulted in approximately 2,600 triggered MS/MS scans with over 70% of triggered scans resulting in peptide spectral matches to the Human UniProt manually annotated database. Roughly 83% of PSMs were derived from unique peptides and resulted in over 600 protein IDs per run. The results from all shotgun proteomic analyses is shown in Table 1 and Figure 2A demonstrates the high overlap in peptide identifications with SequestHT between 3 injected samples.

**Table 1.**
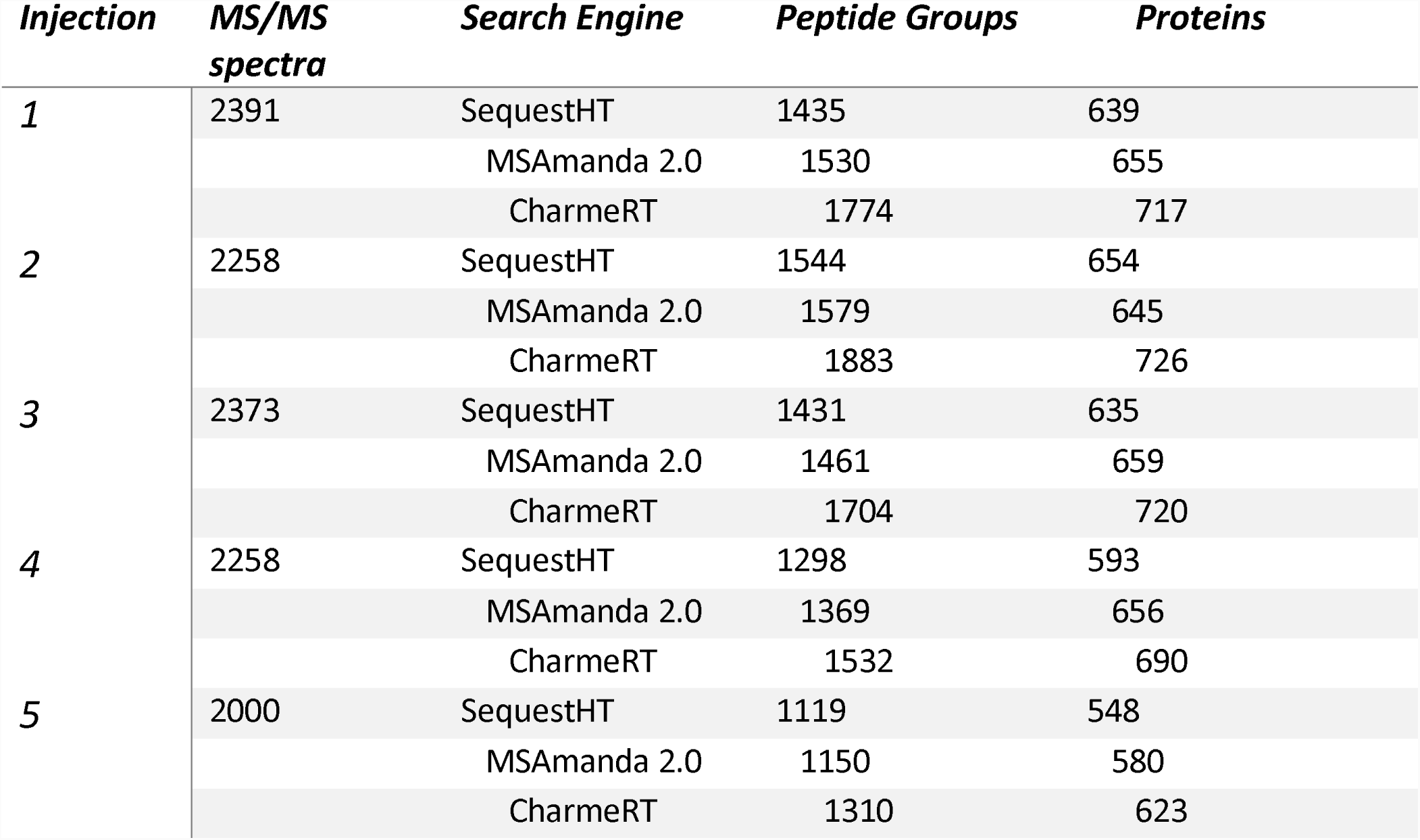
An overview of the results obtained from the Hela peptide injections using different search engines.

| ***Injection*** | ***MS/MS spectra*** | ***Search Engine*** | ***Peptide Groups*** | ***Proteins*** |
| --- | --- | --- | --- | --- |
| **1** | 2391 | SequestHT | 1435 | 639 |
|  |  | MSAmanda 2.0 | 1530 | 655 |
|  |  | CharmeRT | 1774 | 717 |
| **2** | 2258 | SequestHT | 1544 | 654 |
|  |  | MSAmanda 2.0 | 1579 | 645 |
|  |  | CharmeRT | 1883 | 726 |
| **3** | 2373 | SequestHT | 1431 | 635 |
|  |  | MSAmanda 2.0 | 1461 | 659 |
|  |  | CharmeRT | 1704 | 720 |
| **4** | 2258 | SequestHT | 1298 | 593 |
|  |  | MSAmanda 2.0 | 1369 | 656 |
|  |  | CharmeRT | 1532 | 690 |
| **5** | 2000 | SequestHT | 1119 | 548 |
|  |  | MSAmanda 2.0 | 1150 | 580 |
|  |  | CharmeRT | 1310 | 623 |

**Figure 2.**
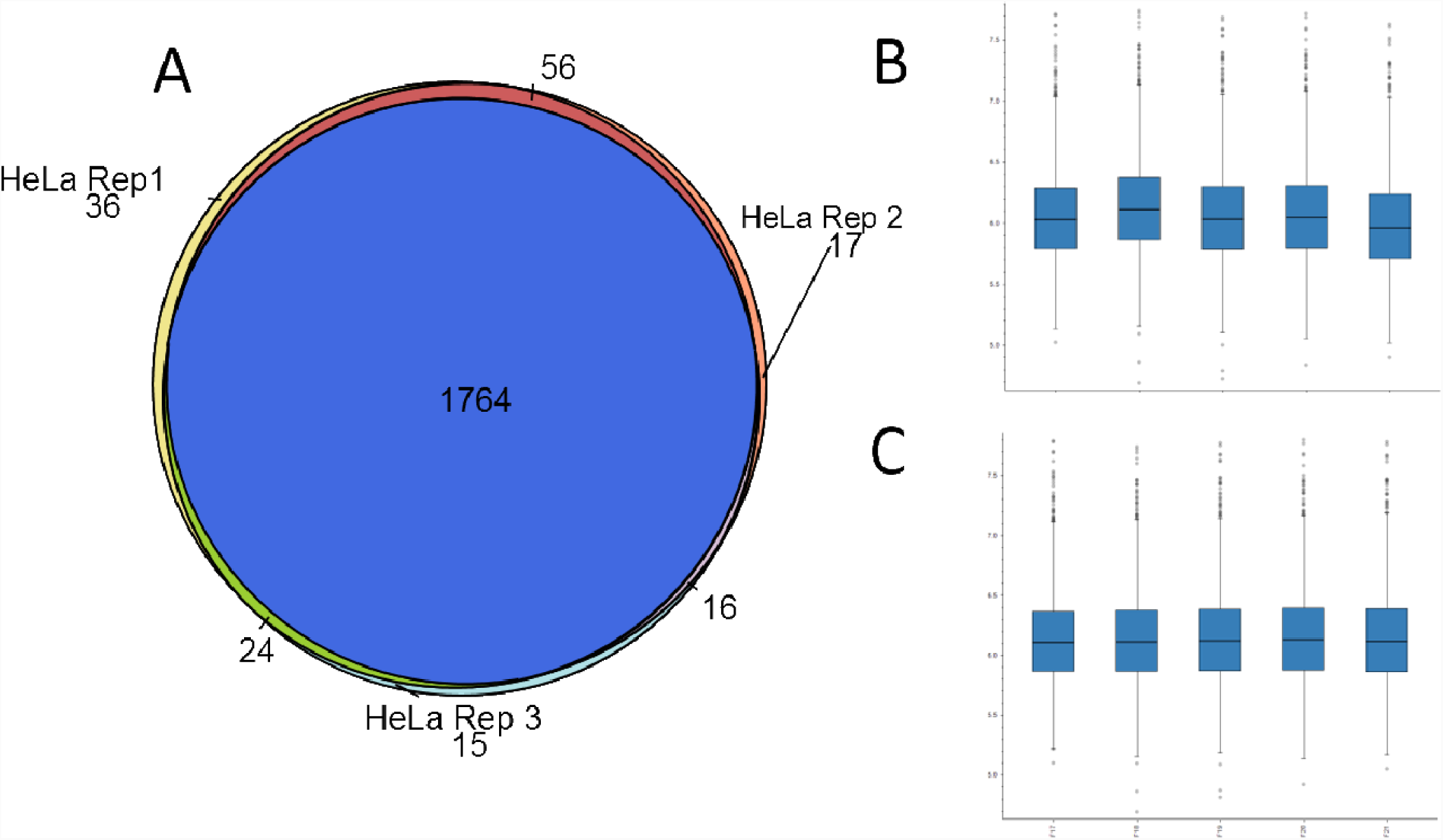
**A)** Unique peptide groups and overlap between 3 technical replicates of Hela digest. B) log transformed loading plots between 5 Hela technical replicates. C) loading plots as in B after normalizing on all peptide averages.

### Second Search with CharmeRT

A recent update to the MSAmanda 2.0 search engine allows the use of the CharmeRT workflow. CharmeRT permits the second search of an MS/MS mass window and precursor coisolation window. Due to the short elution gradient of the complex Hela lysate sample we anticipated high coisolation interference. Searches with and without CharmeRT were performed to test the efficacy of such an approach. Results of the CharmeRT second search are displayed in Table 1. CharmeRT provided an increase of 15.7% in peptide groups and an 8.7% increase in total protein groups over MSAmanda search alone.

### Feature based analysis of Hela lysate

Relative quantification through MS1 based chromatographic feature analysis is becoming common in proteomics applications. MaxQuant lFQ^19^, OpenMS^20^ and more recent algorithms such as IonStar^5^, Minora and apQuant^21^ allow the addition of MS1 feature based relative quantification to be added to nearly any proteomics workflow. Five identical Hela separations were analyzed with the Minora engine in Proteome Discoverer 2.2. The number of unique features per file was very consistent among identical runs (shown in Supplemental table 1), with an average of 19719 ± 268.7 features per file. Both intensity-based abundance estimates per file and peak area-based metrics were tested and demonstrated similar results. Normalization methods used in lFQ were found to be compatible with the CE-MS data as shown in Figure 2B and 2C.

### Targeted quantification in complex digest background

To test the reproducibility of the ZipChip system in peptide elution time, a 400fmol solution of PRTC was spiked into a 200ng/μl Hela lysate background. The 12-minute separation method described above was altered by allowing MS1 scans followed by 4 PRMs on target PRTC peptides. As shown in Figure 3, repeated injections of the same sample loading time exhibited only slight variations in elution pattern. The maximum time variation across runs is shown and approximated a 16 second shift in apex peak elution time. When compared to a recent analysis of repeated runs of a BSA digest standard on 4 different nlC systems^22^ the zCE-MS separation appears comparable.

**Figure 3.**
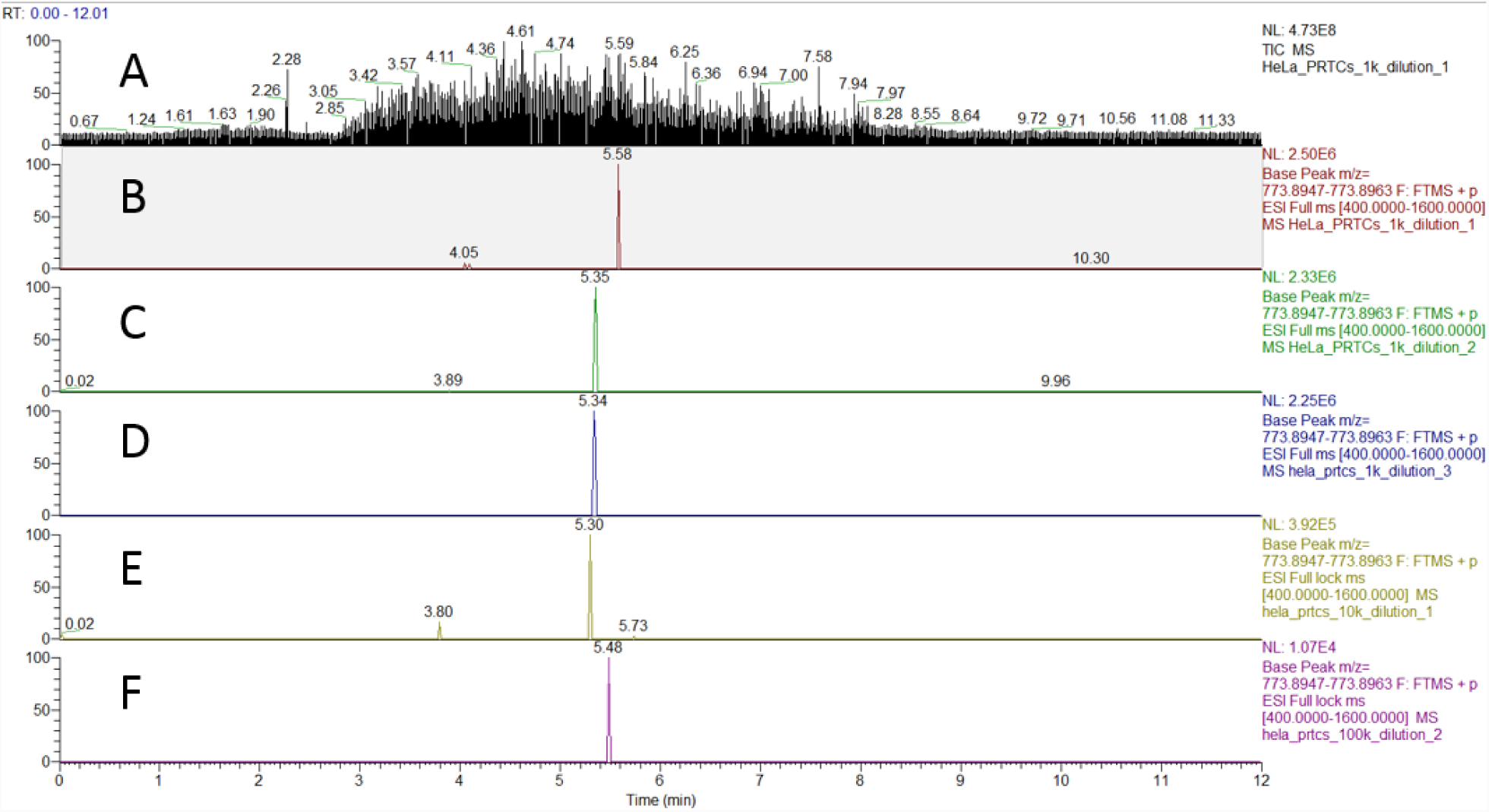
**A** The TIC of the Hela/PRTC background. **B-D** Extracted ion chromatogram from MS1 for the PRTC peptide of identical dilutions. E-F. Extracted ion chromatograms of the same compound at 10x and 100x dilution from above. F represents the lowest dilution of the PRTC peptide that could be detected.

A serial dilution of the PRTC in the same Hela background was also performed to estimate the detection limits within a complex background. A 4fmol injection approximated the limits of detection for the full range MS1 scan, as a 10x dilution from this level did not report signal within the 1ppm MS1 tolerance required. PRM scans were performed on 4 peptides within the mixture using a 100ms fill time. These targeted scans did not provide accurate detection of the injection of the 400attomol PRTC peptides in the 200ng/μl Hela digest solution, suggesting that the limit of detection falls between these marks in a complex lysate background. Clearly, further analysis is required to accurately determine the detection limits of this CE-MS.

Timed or triggered PRM analysis could yield significant improvements in the results by allowing increased ion injection times. However, we feel the largest contribution could be the ability to directly move a single ZipChip system from an Orbitrap-based discovery system to targeted analysis with a triple quadrupole instrument. Automated transfer of target peptide retention times and ease of physical transfer of the ZipChip system could allow both discovery and large cohort validation experiments to be completed in a single day.

## Conclusions

We have evaluated the use of a commercially available capillary electrophoresis system for shotgun proteomics applications. We conclude that rapid separation is an attractive alternative for many proteomics applications where the identification and relative quantification of hundreds of proteins would be a valuable output. While the detection limits of the system are somewhat confounded by questions regarding the amount of peptide loaded into each separation, we conclude that the retention time and reproducibility appears equivalent to traditional nLC-MS techniques. Further analysis will be required to determine if high resolution offline fractionation followed by zCE-MS would provide comparable results to nLC-MS workflows.

## Acknowledgements

We would like to thank Dr. Gary Paul and Mike Goodwin for access to the ZipChip interface and for assistance in operating the interface. Dr. Bob Cole, Dr. Simion Kreimer, and Robert O’Meally of the JHU Proteomics Center provided invaluable discussions for the design and execution of these studies.

## References

(1) Zhang, Y.; Fonslow, B. R.; Shan, B.; Baek, M. C.; Yates, J. R. Protein Analysis by Shotgun/Bottom-up Proteomics. Chemical Reviews. 2013. https://doi.org/10.1021/cr3003533.

(2) Shishkova, E.; Hebert, A. S.; Coon, J. J. Now, More Than Ever, Proteomics Needs Better Chromatography. Cell Systems. 2016. https://doi.org/10.1016/j.cels.2016.10.007.

(3) Kelstrup, C. D.; Bekker-Jensen, D. B.; Arrey, T. N.; Hogrebe, A.; Harder, A.; Olsen, J. V. Performance Evaluation of the Q Exactive HF-X for Shotgun Proteomics. J. Proteome Res. 2018. https://doi.org/10.1021/acs.jproteome.7b00602.

(4) Mertins, P.; Tang, L. C.; Krug, K.; Clark, D. J.; Gritsenko, M. A.; Chen, L.; Clauser, K. R.; Clauss, T. R.; Shah, P.; Gillette, M. A.; et al. Reproducible Workflow for Multiplexed Deep-Scale Proteome and Phosphoproteome Analysis of Tumor Tissues by Liquid Chromatography-Mass Spectrometry. Nat. Protoc. 2018. https://doi.org/10.1038/s41596-018-0006-9.

(5) Shen, X.; Shen, S.; Li, J.; Hu, Q.; Nie, L.; Tu, C.; Wang, X.; Orsburn, B.; Wang, J.; Qu, J. An IonStar Experimental Strategy for MS1 Ion Current-Based Quantification Using Ultrahigh-Field Orbitrap: Reproducible, In-Depth, and Accurate Protein Measurement in Large Cohorts. J. Proteome Res. 2017. https://doi.org/10.1021/acs.jproteome.7b00061.

(6) Shen, S.; An, B.; Wang, X.; Hilchey, S. P.; Li, J.; Cao, J.; Tian, Y.; Hu, C.; Jin, L.; Ng, A.; et al. Surfactant Cocktail-Aided Extraction/Precipitation/On-Pellet Digestion Strategy Enables Efficient and Reproducible Sample Preparation for Large-Scale Quantitative Proteomics. Anal. Chem. 2018. https://doi.org/10.1021/acs.analchem.8b02172.

(7) Shen, X.; Shen, S.; Li, J.; Hu, Q.; Nie, L.; Tu, C.; Wang, X.; Poulsen, D. J.; Orsburn, B. C.; Wang, J.; et al. IonStar Enables High-Precision, Low-Missing-Data Proteomics Quantification in Large Biological Cohorts. Proc. Natl. Acad. Sci. 2018. https://doi.org/10.1073/pnas.1800541115.

(8) Bekker-Jensen, D. B.; Kelstrup, C. D.; Batth, T. S.; Larsen, S. C.; Haldrup, C.; Bramsen, J. B.; Sorensen, K. D.; Hoyer, S.; Orntoft, T. F.; Andersen, C. L.; et al. An Optimized Shotgun Strategy for the Rapid Generation of Comprehensive Human Proteomes. Cell Syst. 2017, 4 (6), 587-+. https://doi.org/10.1016/j.cels.2017.05.009.

(9) Sarg, B.; Faserl, K.; Kremser, L.; Halfinger, B.; Sebastiano, R.; Lindner, H. H. Comparing and Combining Capillary Electrophoresis Electrospray Ionization Mass Spectrometry and Nano-Liquid Chromatography Electrospray Ionization Mass Spectrometry for the Characterization of Post-Translationally Modified Histones. Mol. Cell. Proteomics 2013. https://doi.org/10.1074/mcp.M112.024109.

(10) Khatri, K.; Klein, J. A.; Haserick, J. R.; Leon, D. R.; Costello, C. E.; McComb, M. E.; Zaia, J. Microfluidic Capillary Electrophoresis-Mass Spectrometry for Analysis of Monosaccharides, Oligosaccharides, and Glycopeptides. Anal. Chem. 2017. https://doi.org/10.1021/acs.analchem.7b00875.

(11) Wojcik, R.; Li, Y.; MacCoss, M. J.; Dovichi, N. J. Capillary Electrophoresis with Orbitrap-Velos Mass Spectrometry Detection. Talanta 2012. https://doi.org/10.1016/j.talanta.2011.10.048.

(12) Scheltema, R. A.; Hauschild, J.-P.; Lange, O.; Hornburg, D.; Denisov, E.; Damoc, E.; Kuehn, A.; Makarov, A.; Mann, M. The Q Exactive HF, a Benchtop Mass Spectrometer with a Pre-Filter, High-Performance Quadrupole and an Ultra-High-Field Orbitrap Analyzer. Mol. Cell. Proteomics 2014. https://doi.org/10.1074/mcp.M114.043489.

(13) Kelstrup, C. D.; Jersie-Christensen, R. R.; Batth, T. S.; Arrey, T. N.; Kuehn, A.; Kellmann, M.; Olsen, J. V. Rapid and Deep Proteomes by Faster Sequencing on a Benchtop Quadrupole Ultra-High-Field Orbitrap Mass Spectrometer. J. Proteome Res. 2014. https://doi.org/10.1021/pr500985w.

(14) Mellors, J. S.; Gorbounov, V.; Ramsey, R. S.; Ramsey, J. M. Fully Integrated Glass Microfluidic Device for Performing High-Efficiency Capillary Electrophoresis and Electrospray Ionization Mass Spectrometry. Anal. Chem. 2008. https://doi.org/10.1021/ac800428w.

(15) Wang, Y.; Feng, P.; Sosic, Z.; Zang, L. Monitoring Glycosylation Profile and Protein Titer in Cell Culture Samples Using ZipChip CE-MS. J. Anal. Bioanal. Tech. 2017. https://doi.org/10.4172/2155-9872.1000359.

(16) Eng, J. An Approach to Correlate Tandem Mass Spectral Data of Peptides with Amino Acid Sequences in a Protein Database. J. Am. Soc. Mass Spectrom. 1994. https://doi.org/doi:10.1016/1044-0305(94)80016-2.

(17) Dorfer, V.; Pichler, P.; Stranzl, T.; Stadlmann, J.; Taus, T.; Winkler, S.; Mechtler, K. MS Amanda, a Universal Identification Algorithm Optimized for High Accuracy Tandem Mass Spectra. J. Proteome Res. 2014. https://doi.org/10.1021/pr500202e.

(18) Dorfer, V.; Maltsev, S.; Winkler, S.; Mechtler, K. CharmeRT: Boosting Peptide Identifications by Chimeric Spectra Identification and Retention Time Prediction. J. Proteome Res. 2018. https://doi.org/10.1021/acs.jproteome.7b00836.

(19) Cox, J.; Hein, M. Y.; Luber, C. A.; Paron, I.; Nagaraj, N.; Mann, M. Accurate Proteome-Wide Label-Free Quantification by Delayed Normalization and Maximal Peptide Ratio Extraction, Termed MaxLFQ. Mol. Cell. Proteomics 2014. https://doi.org/10.1074/mcp.M113.031591.

(20) Weisser, H.; Nahnsen, S.; Grossmann, J.; Nilse, L.; Quandt, A.; Brauer, H.; Sturm, M.; Kenar, E.; Kohlbacher, O.; Aebersold, R.; et al. An Automated Pipeline for High-Throughput Label-Free Quantitative Proteomics. J. Proteome Res. 2013. https://doi.org/10.1021/pr300992u.

(21) Doblmann, J.; Dusberger, F.; Imre, R.; Hudecz, O.; Stanek, F.; Mechtler, K.; DUrnberger, G. ApQuant: Accurate Label-Free Quantification by Quality Filtering. J. Proteome Res. 2018. https://doi.org/10.1021/acs.jproteome.8b00113.

(22) Liu, Q.; Cobb, J. S.; Johnson, J. L.; Wang, Q.; Agar, J. N. Performance Comparisons of Nano-LC Systems, Electrospray Sources and LC-MS-MS Platforms. J. Chromatogr. Sci. 2014. https://doi.org/10.1093/chromsci/bms255.

